# Prioritization of decisions by importance and difficulty in human planning

**DOI:** 10.1101/2024.10.19.619044

**Authors:** Fatmaalzahraa A Aboalasaad, Leonie Lambertz, Chiara Mastrogiuseppe, Rubén Moreno-Bote

## Abstract

In our everyday lives, we continually need to commit to courses of future action even when direct feedback is not available. The expected reward of an action depends not only on a single decision but on a sequence of interdependent choices that will happen in the future. Admittedly, most decisions we make concern sequences of actions as opposed to single-step choices. To make these decisions, we rely on our ability to plan and forecast the potential outcomes of sequential decisions. However, how humans plan under novel real-world scenarios remains poorly understood. We developed a novel task to investigate how humans evaluate options and prioritize their decisions during planning in realistic situations. In each trial, a written planning scenario (for example, “plan your birthday party”) was followed by 9 pictures divided into three categories (e.g., 3 birthday cakes, 3 party locations, and 3 decorations). Participants were asked to choose one option from each category to create the best possible plan - the one with the highest subjective value – while both their response and gaze were tracked. With each option possibly having a different subjective value according to the other selected options, the required planning process resembles the navigation of an internal decision tree, whose complexity grows exponentially with the number of choices and future outcomes considered. Our results show that participants gather information at all levels of the decision trees, as suggested by their gaze-switching behavior. In addition, based on their assessments of importance and difficulty, we find that participants generally choose first what they report to be the most important and easiest category and then the least important and most difficult one. Overall, our task provides a novel means to study planning behavior in realistic, multi-alternative situations, as participants can freely navigate through all levels of the decision tree by subjectively evaluating potential scenarios through internal sampling and imagination.

## Introduction

Imagine your birthday is approaching, and you are planning to host a party. You will need to make decisions about several factors, such as the venue, menu, and guest list. Each choice involves considering different options and combinations, requiring you to commit to future actions without immediate feedback (Sims et al., 2013). Importantly, the expected reward for the final plan is not dependent on a single action but on an interdependent sequence of decided future actions (Jain et al., 2019; Sutton & Barto, 1998). To accomplish these plans, you rely on your ability to plan and predict the possible outcomes of successive decisions (Keramati et al., 2016; Miller & Cohen, 2001). Humans and animals are capable of solving complex tasks by relying on their ability to plan, i.e., by extracting the sequence of actions they should perform to achieve their goals. In this context, thinking about the future reward and the actions one could take to achieve a goal can be likened to the exploration of a decision tree whose complexity grows exponentially with the horizon considered (Huys et al., 2015; Sutton & Barto, 1998; Mastrogiuseppe & Moreno-Bote, 2022).

In essence, planning can be defined as a sequence of interdependent actions that have yet to be executed (Balleine & Dickinson, 1998) and hence, it involves two essential calculations: predicting the outcome of actions and evaluating their utility. This process relies on an internal search for information based on past experiences (Hunt et al., 2021), which allows for better predictions about the future (Daw & Dayan, 2014; Hunt et al., 2021; Shohamy & Daw, 2015; Shadlen & Shohamy, 2016). Having planning abilities is crucial to adapt to changes in the environment that cannot be anticipated and allow agents to behave flexibly. However, planning comes at a high cost, requiring many resources to be run (memory, and computational power, although time is arguably the major limitation) (Mattar & Lengyel, 2022). Indeed, the estimation of the potential rewards associated with a specific course of action within a decision tree relies on the agent’s mental simulation of the environment using its internal model. This internal model is essentially a learned, subjective representation of the environment based on past experiences (Ho et al., 2022; Pezzulo et al., 2019; C. E. Sezener et al., 2019). In theory, the agent can simulate every conceivable sequence of actions and their resulting outcomes to deduce the most advantageous plan leading to the highest utility outcome (Sutton & Barto, 1998). However, the practical application of such an exhaustive search within the decision tree is unfeasible because the number of potential action sequences grows exponentially with each additional step into the future, rendering it too complex and resource-intensive to be practically executed (Huys et al., 2012; Mastrogiuseppe & Moreno-Bote, 2022). The computational load of such can be minimized by employing habitual behavior in some parts of the decision tree, which uses previous, well-established knowledge of the value of states in the future and how to act on those states (Otto et al., 2013, Balleine & O’Doherty, 2010; Dolan & Dayan, 2013; Keramati et al., 2011, 2016).

Numerous studies have examined human strategies for planning. Simon (1973) argued that expert chess players mentally compute a maximum of about 100 possible moves but rarely explore more than ten moves in depth. This limited depth itself can be seen as a heuristic, as it reflects the necessity of approximations when the full decision tree cannot be fully explored. Similarly, Keramati et al. (2016) observed that humans employ heuristics such as pruning the decision tree and generating subgoals while exploring it. These findings highlight how humans largely rely on heuristics to manage planning complexity. Further psychological studies show that humans reduce planning costs through prioritizing and pruning action sequences (Daw et al., 2005; Gläscher et al., 2010; Ramkumar et al., 2016). Many studies showing different strategies used during planning also include the use of an internal representation of the environment to predict outcomes. This is consistent with the principles of model-based reinforcement learning, where the agent relies on a simulated model to predict outcomes (Dolan & Dayan, 2013). Alternatively, other approaches are based on trial and error, where actions are performed iteratively to obtain a reward (Cushman & Morris, 2015).

Recent studies have advanced our understanding of human planning by designing experiments in which participants can control certain factors, such as the number of options to consider in the first place, while other factors are constrained by the experimental design, like the position of the options in the decision tree. For instance, hierarchical tasks for perceptual decisions have been developed in which subjects can uncover a hidden target by integrating stochastic cues to moving points at the nodes of the decision tree (Zylberberg, 2021). The study demonstrates that participants adeptly navigate decision-making hierarchies through strategic planning and reasoning by categorizing decisions, filtering unreliable sensory information, and using their confidence to identify potential errors. Keramati and colleagues (2016) found that the degree of planning depth plays a crucial role in influencing the balance between habitual and goal-directed behaviors. A more superficial planning approach was associated with an increase in habitual behavior, while deeper planning strategies were associated with an increase in goal-directed behavior. Adaptive integration of habits into the planning process was found, and this integration was influenced by variables such as the rarity of transitions and the presence or absence of rewards (Keramati et al., 2016). These results are in line with another study investigating the effect of expertise in planning. By examining a complex board game and using a cognitive model based on a heuristic search to capture human behavior and predict decisions, reaction times, and eye movements, the results show robust evidence for increased depth of planning with expertise (van Opheusden et al., 2023). Another study on Monte Carlo Tree Search (MCTS) introduces a new perspective by quantifying the computational value based on statistics of possible future action consequences and decision-making that takes into account the expected impact on the quality of the chosen action (Sezener & Dayan, 2020).

Despite the progress in our understanding of planning, a full picture of the strategies that humans use is incomplete. One limiting factor is that most tasks developed to study planning so far lack realism, as participants must cope with hypothetical, abstract rather than real-life scenarios. As Schulz et al. (2018) emphasize, to make real planning comprehensively, one must not only evaluate the consequences of decisions but also anticipate the future motivational state of the planning agent (Schulz et al., 2018). Further, most previous tasks do not consider the inherent value of choices because they overlook the interdependence (i.e., sequence dependence) of values that are crucial for planning in real life. Rather than using fixed reward amounts on each visited state independent of previously obtained ones, real-life scenarios incorporate subjective values that are strongly dependent on the previously visited states and values. Further, when planning, individuals draw on past experiences to predict the possible outcomes of unknown situations (Mattar & Daw, 2018; Shohamy & Daw, 2015). While decisions based on subjective values, so-called value-based decisions, have been extensively studied in single-step tasks, their exploration in the planning domain is still limited (Brus et al., 2021; Callaway et al., 2022). Especially when sequential decisions are involved, the value of an option varies not only with the environment and the individual’s motivational state but also with other decisions (Krusche et al., 2018). An additional critical difficulty in real-life planning is that there is no initial reference point or predetermined hierarchical structure for the decision tree, but it rather has to be built on the fly. Previous tasks are based on board games or pre-defined trees, and thus the order in which decisions ought to be performed is highly constrained, contrary to many real-life scenarios.

To address the above issues, we devised a novel task where human participants were presented with a realistic planning task. We asked participants to make a plan in scenarios like designing a birthday party or planning their Sunday breakfast. Our main question is how humans build a hierarchy of decisions on the fly in a planning task where no predefined, a priori hierarchical structure is provided. By tracking their gaze, we find that participants explore all levels of the decision tree. Importantly, when participants begin to decide, they typically select items they subjectively consider to be the easiest and most important choices, then proceed to address the remaining choices. Our results shed new light on the calculations underlying human planning.

## Results

### Gaze-switching behavior

We investigated how human participants (N=28) prioritize information and choices in a novel task where the order of information gathering, and choices are not pre-established (Fig. 1). In the task, participants used a mouse to report their choices while their eye movements were recorded. Each trial began with a written planning scenario displayed for 6 seconds (e.g., “Plan your birthday party”). A fixation cross appeared for 1 second, followed by the presentation of 9 images divided into 3 categories (e.g., 3 birthday cakes, 3 party locations, and 3 decorations) for 6s, displayed concentrically around the fixation cross. These categories were distinguishable by randomly assigned colored frames around the images. In each scenario, participants have to create the most desirable plan – the one with the highest subjective value - by selecting one option from each category, in any order and using as much time as they want, with a minimum time of 5s. Once a choice was made, it could not be undone and options of the corresponding category became unavailable, as indicated by the disappearance of the frames around them. Following their choices, participants were asked to express their confidence in the choice of the selected options and their assessment of the importance and difficulty of the categories using a rating scale of 0 to 10.

**Figure 1.**
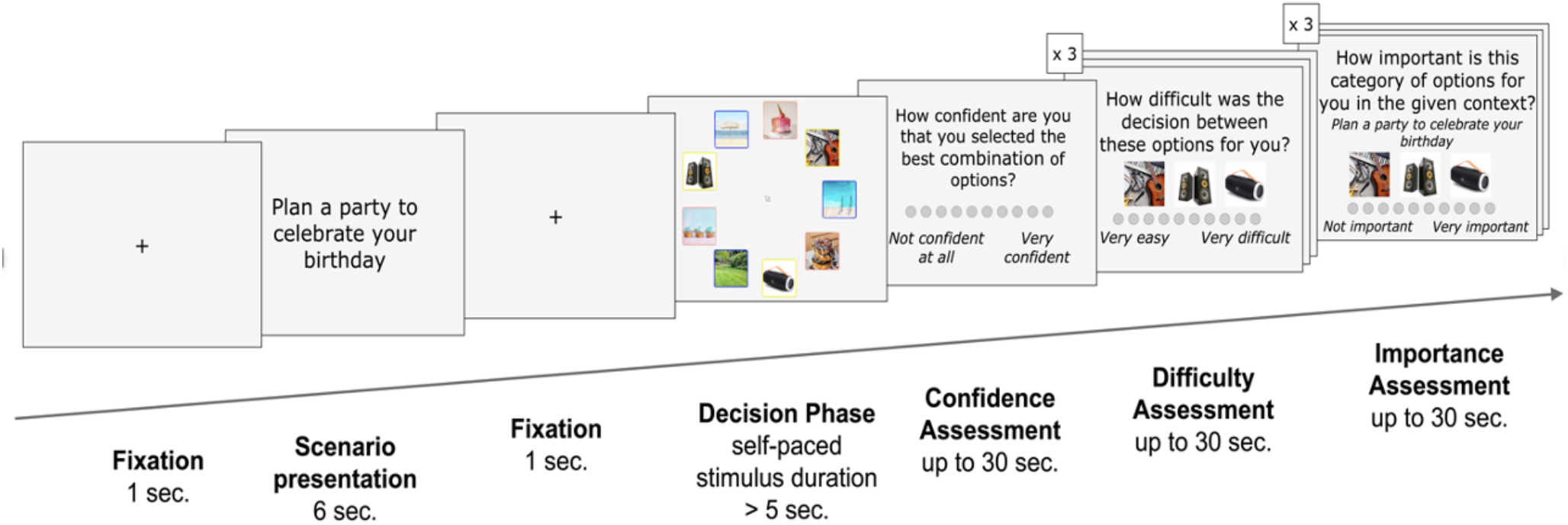
Schematic of a single trial of the context-dependent planning task. The trial begins with the presentation of a fixation cross (fixation not enforced), followed by a 6-second display of written context. Subsequently, a second fixation cross is presented. After this, 9 pictures divided into 3 different categories were displayed concentrically around a fixation cross. Participants were asked to choose one picture from each category by a mouse click. No order was imposed: participants freely chose the order of the categories in which they would choose an image. Then participants were asked about their confidence in choosing the best combination of options, followed by the importance of each category and the choice difficulty. A total of 45 different planning scenarios were tested (Methods).

We find participants executing a complex pattern of eye movements to gather information about the images and make three consecutive choices (Fig. 2a, b), typically wandering over images within a single or several categories before making the first choice. We studied the average time to make the first, second, and third choices. Specifically, RT1 refers to the time from the onset of the stimulus to the selection of the first option from the first category, RT2 is the time taken to make the second choice, starting immediately after the first choice is made and lasting until the second choice is selected, and RT3 is the time taken to make the third choice, starting immediately after the second choice is made and lasting until the third choice is selected. Our results indicate that the pace at which they make the decisions is slow both overall (with mean ± std = 21.23 ± 8.80s) and at single choice level (mean ± std are RT1: 9.61 ± 6.01s, RT2: 6.07 ± 4.30s, and RT3: 5.54 ± 3.95s) (Fig. 2c) when compared to literature (Fontanesi et al., 2019). The longer reaction time for the first choice might imply that participants first engage in a sampling phase where information is gathered and the decision tree is built, as also suggested by further evidence below. Participants directly look (i.e., for at least 30ms) at all the images in most of the trials (88%, Fig. 2d). In only 8% of the trials, the first choice was made before all images across the three categories were sampled, likely due to strong individual preferences or habituation. However, the vast majority of trials demonstrate that decisions were made after thoroughly evaluating all available options. At the time of the three choices, participants did not consistently look directly or uniquely at the selected picture, often looking at options that were different from their reported decision. On average (mean ± std), participants looked at the chosen picture only 36 ± 13% of the time during the first choice, while 24 ± 16% for the second choice, and 36 ± 13% for the third choice. We also observe that participants look back to a category whose preferred item has already been selected when it is time to choose from the second category (fraction of revisits after the first choice over all trials, mean ± std = 9.46% ± 7.85). This observation suggests that participants may need to reevaluate options based on their previous choices and reconsider the choices yet to be made. Moreover, participants not only shift their gaze within a category to compare options but also repeatedly switch between categories (see Fig. 2b).

**Figure 2.**
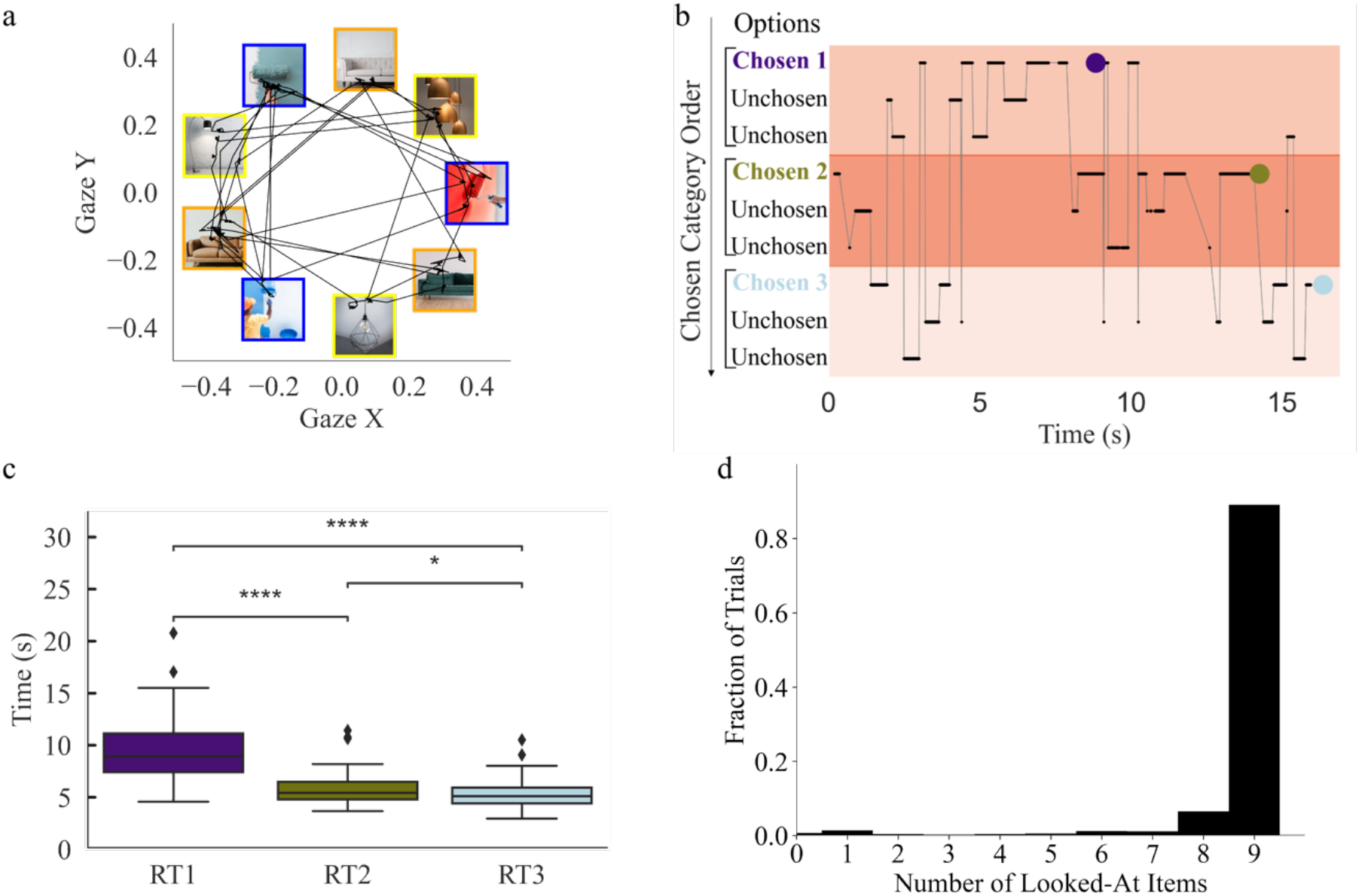
Participants typically sample from all categories and their respective images to develop an informed plan. (**a**) Eye position on normalized screen coordinates for one participant in one typical trial. In this trial, the participant must plan the interior design of their living room, with categories of paint color, sofa, and lamp. (**b**) Gaze position over time in the same trial. Categories are displayed from the top to the bottom according to the decision order of the trial, that is, the category of the first chosen option is displayed at the top, and so on. Purple, green, and cyan dots mark the selection of the images for the first, second, and third choices, respectively. (**c**) The reaction time (RT) for each choice. Colors as in (b). Boxplots were calculated over the means of all participants. Results of pairwise comparisons are displayed based on adjusted p-values (‘*’: P < 0.05, ‘***’: P < 0.001,’****’: P < 0.0001). (**d**) Distribution of the fraction of trials in which participants only look at none, 1, 2, 3, and so on, options in the trial.

### Decision tree structure formation and choice prioritization

An important question is how participants dynamically create their plan, in particular how they mentally navigate the different categories and the images within each category to form a mental tree. Although it is not possible to determine the internal states of the participants, we use their gaze behavior, choices, and timings to infer the potential internal dynamics that underlie planning in this unconstrained planning problem. To investigate whether there is a general way of assembling the levels in the decision tree (that is, what category is chosen from first, and so on), we analyzed the self-reports of the importance and choice difficulty of each of the three categories presented within a trial (Supplementary Fig. S1b).

We find that participants begin by choosing what they report to be the easiest category, followed by the second most difficult, and finally the most difficult category (Supplementary Fig. S1b left panel). We conducted a within-participant analysis ANOVA on the mean perceived difficulty across all participants between categories. To address the problem of subjectivity, in each trial, we call C1, C2, and C3 respectively the first, second, and chosen categories. The analysis revealed a significant difference between all the categories (F = 39.6, P = 0.009). We also found a significant difference between pairs of categories, as quantified by the post-hoc pairwise comparison with Bonferroni correction (between C1 and C2: t = -5.275 e+00, P = 4.367 e-05; C2 and C3: t = -5.118 e+00, P = 6.652 e-05; C1 and C3: t = - 7.232 e+00, P = 2.661 e-07).

Likewise, the reported importance of the categories also correlates with the choice order (Supplementary Fig. S1b right panel; within-participants ANOVA on the mean scores of all participants to examine whether perceived importance differed between the three choices, F = 26.2, P = 1.1e-08). Specifically, participants typically begin with the category they identified as most important, followed by the second most important category, and ultimately conclude with the least important category (post-hoc pairwise comparison with Bonferroni correction; between first and second choices: t = 2.695 e+00, P = 3.590 e-02; between second and third choice: t = 4.276 e+00, P = 6.380 e-04; and between first and third choices: t = 6.899 e+00, P = 6.167 e-07).

The result of our multinomial logistic regression is that participants tend to start with the easiest and most important category held at the individual level when both category importance and difficulty are used as regressors to predict the order of the category choice (see Eqs. 1 and 2 in the Methods). Specifically, the parameters of the logistic regression measure how the regressors influence the likelihood of choosing that category second versus first, and third versus first, respectively. Therefore, a negative value of, e.g., parameter *β*_12_ would mean that increasing the value of the importance regressor decreases the probability of choosing that category as the second in comparison with choosing it first. This is indeed what we find for the effects of importance in both log-odds, while the opposite effects were observed for difficulty (t-test on average parameters across individuals; *β*_02_= -0.08,t = -0.37, P = 0.71, *β*_12_ = -0.034, t = -1.5, P = 0.13, *β*_22_ = 0.107,t = 4.32, P = 1.88 e-04, *β*_03_ =0.01,, t = -0.07, P = 0.94, *β*_13_ = -0.098,t = -4.6, P = 8.78 e-5, *β*_23_ = 0.182, t = 5.65, P = 5.27 e-06). Therefore, at the group level, the higher the importance and the lower the difficulty of a category the more likely it is that it is chosen before others.

In conclusion, participants tend to either simplify their decision tree by prioritizing the simplest decision first or by focusing first on the most important category to make the first choice within that category, revealing a general strategy to build the decision tree. We remark that our experiment allowed us to address this question as the hierarchical structure of the tree is not experimentally enforced.

Finally, we find that participants frequently shift their gaze across images belonging to different categories. In fact, across participants, the across-category gaze switches (moving eyes from an image in one category to an image in another category) are more frequent than the within-category gaze switches (moving eyes between images belonging to the same category), and this was largely consistent across the three different choices (Supplementary Fig.S1a). However, this result may be influenced by the geometrical organization of the images, with images belonging to different categories being displayed nearby. Indeed, we observe that the probability of switching gaze from one image to one of the two nearby images was almost twice as high as switching to one of the six distant images (0.199 vs 0.102; probabilities are normalized by the number of images considered).

### Looking time across categories as a function of importance and difficulty

Further evidence of the importance and difficulty-based structure of the decision tree is the observation that participants tend to focus primarily on the selected category rather than the other two, across all three choices (Supplementary Fig. S1c). We calculated the mean fraction of looking time at the first, second, and third chosen categories (C1, C2, and C3) for each participant within each of the three choice periods. We find that from the onset of stimuli to the first choice, C1 is observed for the longest fraction of time (Supplementary Fig. S1c; within-participant analysis ANOVA, F = 92.7, P = 3.4 e-18). During the time taken to make the second choice, it is C2 the one that is most gazed at (within-participant analysis ANOVA, F = 171.14, P = 4.3 e-24). Finally, during the time taken to make the third choice, it is C3 the category mostly looked at (within-participant analysis ANOVA, F = 309.76, P = 2.6 e-30). For the second and third choices, the fraction of time that the first chosen category C1 is gazed at is very small (0.11 and 0.09, respectively). This suggests that, while participants may observe all categories throughout the decision-making process, they tend to focus on a single category after the initial sampling period (early during the first choice). This indicates that the category order is likely, at least partially, determined during the early stages of planning.

Participants tend to spend a large fraction of time focusing on the first chosen category (Supplementary Fig. S1c, left panel). Relevantly, we find this result not to be restricted to the cases where the first chosen category is the most important one, as it would be expected in cases where importance drives the selection process, but also when C1 is the category reported as the easiest. This is consistent with the earlier idea that participants engage in extensive sampling of all options to build their decision tree. Therefore, we studied how the amount of time that participants looked at corresponding to the time spent to choose each category for each trial, denoted RT, simultaneously depended on importance and difficulty. We regressed the log (RT) for each participant based on the importance and difficulty reported by the participant across trials treated as different regressors and analyzed the effects at the group level (Eq. 3 in the Methods). We first modeled the data for each participant, then calculated the average weights of the participants and performed a t-test to determine whether the average weights were significantly different from zero. We find that while both regressors have a positive weight, only difficulty significantly increases the decision time for the first choice (*β*_0_=1.52, with t = 23.16, P = 1 e-300; *β*_11_ = 0.0037 with t = 0.64 and P = 0.53; *β*_12_ =0.054, with t = 11.01 and P = 1.71 e-11). The effect of category importance, although having a significant effect on decision time, was much weaker compared to the effect of choice difficulty (t = -6.2, P = 1.24 e-06). These results do not contradict the above ones regarding the ordering of choices: people typically start with the easiest category, but if the first chosen category happens to be not that easy, then participants need more time to choose among the images that it contains.

Gaze patterns correlate with the confidence level that participants have in each choice. In line with this, we find that participants tend to perform a larger number of gaze switches in those trials where their overall confidence in their choice was lower (average Spearman’s correlations = -0.336; Wilcoxon test, V = 0, P = 3.725 e-09, testing whether the rho’s are negative at the population level).

This suggests that participants not only take into account category importance and difficulty in building the decision tree but the pace at which this is made is strongly modulated by their confidence. Indeed, using a linear model that predicts choice confidence per participant across trials based on importance, difficulty, log(RT), and the number of switches (Eq. 4 in Methods), we find that choice confidence depends on the average importance and difficulty over the three categories presented in each trial, with importance increasing choice confidence while difficulty reducing choice confidence (*β*_0_=12.01, with t = 12.5, P = 8.85 e-13; *β*_11_ = 0.45, t = 6.6 and P = 4.34 e-07; *β*_12_ = -0.83, t = -8.2 and P = 6.61 e-09 ; *β*_13_ = -1.5, t = -5.001 and P = 2.99 e-05, *β*_14_= -0.004, with t = 0.85 and P = 0.4).

### Simplicity and speed can account for why participants tend to begin with the most important category first and/or the easiest category

There could be multiple reasons why people tend to start the planning process by choosing the most important category and/or the easiest. Choosing the most important category indicates that the participants prefer to make the most important decision first, so that the next two, less important decisions can be aligned with it. Choosing from the easiest category first might imply a different strategy, complementary to the previous one: by first making the easiest choice, participants can easily simplify the decision tree, making faster decisions overall. Both strategies seem to obey the same rationale of simplifying the planning process and making it faster. It is conceivable that starting from the least important category could have deleterious consequences, as once arriving at the choice of the more important category a decision might imply the need of undoing a former choice. Similarly, starting with a more difficult category can lead to spending considerable time on a decision that may need to be reversed once easier decisions have been made. As in our task participants can decide where to start from, this degree of freedom seems to reveal a common strategy that serves to efficiently solve complex, realistic planning problems.

## Discussion

We investigated how people prioritize their decision-making processes in the planning process. Our results reveal a clear pattern in which participants consistently prioritize the most important categories when creating decision trees, followed by the less important ones. They also tend to start with the easiest decision, followed by the more difficult ones. The observed behavior might correspond to the innate human tendency to simplify and speed up the planning phase. Our task requires foreseeing future consequences of the choices, very often heavily intertwined so that the optimal choice at one level strongly correlates with the optimal choice at another level. Our results reveal that simple and consistent heuristics are employed in the formation and resolution of decision trees in human planning.

Answering the question of how humans build decision trees requires some innovation in the experimental paradigm. A notable innovation in our methodology is that the structure of the decision tree needed for planning is not given, but it has to be built on the fly by participants. This freedom of choice is a distinguishing feature that sets our experiment apart from other studies on planning (Akam et al., 2015; Callaway et al., 2022; Daw et al., 2011; Keramati et al., 2016; Zylberberg, 2021). Participants were able to choose the combination of scenarios that were most valuable to them personally, which permits the study of the dynamics of the decision tree construction. This aspect introduces a dynamic element that allows us to examine the intricate interplay of individual preferences in the decision-making process.

We also observed that participants often switched their gaze between different categories more frequently than within the same category. This could indicate that they were thinking strategically about the relationships between categories before evaluating options within each category. The research findings emphasize the importance of imagination in human planning, in which the evaluation of possible courses of action involves a mental simulation (Krusche et al., 2018; Nanay, 2016). Additionally, we found that when participants were less confident in their choices, they shifted their gaze more frequently to gather more information. This sheds light on the dynamic relationship between visual attention and decision confidence in the planning and evaluation phases of decision-making. Thus, the finding extends previous work on single-step decisions by demonstrating a comparable effect in sequential value-based choices (Call & Carpenter, 2001; Song et al., 2019; Drugowitsch et al., 2014; Vickers, 2014).

Our goal is to decipher the factors that influence the time required for option evaluation by examining the correlation between the proportion of attention directed to the categories and their respective levels of choice difficulty and importance. We found that participants spent more time on more difficult categories, reflecting the cognitive effort required for tough decisions. This result fits seamlessly with the findings from studies on value-based decision-making and emphasizes the importance of cognitive effort when faced with difficult decisions (Ludwig & Evens, 2017). In contrast, we observed an opposite trend when considering the importance of a category. Participants spent less time selecting a category that they considered important. This inverse relationship suggests that the perceived importance of a category rationalizes the decision-making process and leads to a faster choice. In addition, the amount of time participants spent looking at each category proved to be a valuable metric. By analyzing how long participants focused on each category, we gained insights into the order and hierarchy of decisions, shedding light on how decision trees are built (Cavanagh et al., 2014; Spering, 2022).

While our findings offer valuable insights into human planning, there are some limitations. In each trial of the current study, participants were presented with a set of 9 options, which may not fully capture the complexity of larger planning problems (Mastrogiuseppe & Moreno-Bote, 2022). Nevertheless, the lack of a predetermined decision tree structure and the interdependence of option values contribute to increased task complexity and realism, reflecting the challenges encountered in real-life planning scenarios (Wu et al., 2017; Zylberberg, 2021). Extending the task to larger state spaces proves to be a promising avenue for future research (van Opheusden & Ma, 2019).

All in all, we have developed a novel human planning task that permits exploring planning strategies in somehow realistic scenarios. Participants evaluate options subjectively by drawing on their imagination and memory and have the flexibility to build and move freely through all levels of the decision tree. This approach has allowed us to identify common patterns in decision prioritization (“more important and easier first”) and option evaluation that shed light on common strategies used by individuals in the human planning domain.

## Methods

### Participants

The sample consists of 28 participants. All participants had normal or corrected-to-normal vision. The full length of the experimental session was approximately one hour including 45 trials. Participants were recruited through the CBC participant database. Participation in the study was financially compensated with 10 €. The research project received ethical clearance from the Institutional Commission for Ethical Review of Projects (CIREP) of Pompeu Fabra University. Before participation, the participants were presented with a comprehensive informed consent form outlining the experimental protocol, its duration, participants’ rights, and data confidentiality. After completion of the study, the participants received a debriefing that included the experimenter’s contact information for future inquiries.

### Behavioral task

The task was performed on a computer screen while participants’ gaze positions were recorded. Participants underwent an intake procedure before the experimental session in which they were required to sign the consent form and read the task instructions. During the experiment, they were instructed to sit in a comfortable position on the chair and to move their head as little as possible. Each participant’s calibration profile was created using a 5-point calibration component of PsychoPy. After the calibration process, participants completed a training phase consisting of two trials to familiarize themselves with the task. Each trial began with a written planning scenario for 6 seconds (e.g., “Plan your birthday party.”). Then, a fixation cross was displayed for 1 second. Then, 9 images were displayed, which were divided into 3 categories (e.g., 3 birthday cakes, 3 party locations, and 3 decorations) for 6 seconds. The categories were recognizable by the colored frames around the images. The frame color of each category was randomly assigned. In the given scenario, participants were asked to choose one option from each category to create the best possible plan - the one with the highest subjective value. During the decision phase, images are displayed in a circular arrangement on the screen. The spatial locations of these images form a nine-sided polygon, with none of the locations lying on the vertical or horizontal diameters of the circle. This design choice aims to avoid saliency effects caused by geometric symmetries. Every third image along the circle represents the same ategory. This means that all options within a category are evenly spaced. The possible image locations remain consistent throughout the experiment, allowing participants to become familiar with the category arrangement and potentially reducing visual search time during the decision phase. The specific image locations are randomly assigned to the images in each trial. After the final choice had been made, we asked participants about their choice confidence. Further, participants reported the subjective importance of the categories and choice difficulty. The difficulty, importance, and confidence were rated using a rating scale ranging from 0 to 10. The starting position of the slider on the scale was randomized to avoid response biases.

Upon completion of all trials, a text appeared on the screen signaling successful task completion, and the experimenter entered the room to debrief the participant. To make the choice, participants use the mouse located in the center of the circle containing the images. We have 45 different scenarios divided into 3 different categories as shown in the supplementary Table.ST1. When considering a planning scenario, e.g., planning a birthday party, participants are faced with three different categories, e.g., choosing the place, the menu, and the music. When constructing a decision tree for this scenario, the question arises as to which category should be the starting point for building the tree: should one begin with the place, the food, or the music?

### Stimuli

Visual stimuli were presented on the Asus 27-inch monitor with a 1920×1080 pixel resolution and a frame rate of 60Hz. Participants had to hold onto a computer mouse on a table using their dominant hand. The eye-tracking device Tobii Pro Spectrum recorded gaze data from participants throughout the planning task at 120 Hz. The number of planning scenarios was 45 in the experiment. The scenarios are shown in the supplementary Table.ST1.

## Analysis

### Probability of order of category as a function of importance and difficulty

We applied a multinomial logistic regression model to predict the order in which a category is chosen depending on the reported importance and difficulty of the 3 categories. In this model, the dependent variable represents the chosen order (C1, C2, C3) of a specific category, and the independent variables (regressors) include the importance and difficulty of that category

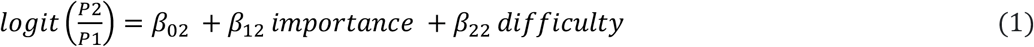

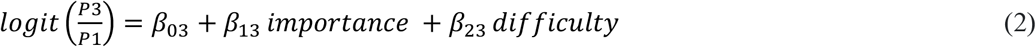

where *P*1 is the probability that category 1 is chosen in the first position, *P*2 is the probability that category 2 is chosen second, and so on. Importance and difficulty represent the ratings given by the participants based on their subjective assessment of the importance and difficulty of each category. We first modeled the data for each participant, then calculated the average weights of the participants and performed a t-test to determine whether the average weights were significantly different from zero.

### Factors that influence the time of choice

We applied a multiple linear regression model to evaluate the effects of importance and difficulty of categories on log-transformed reaction times for each participant.

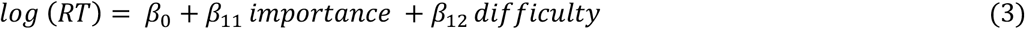

where RT is the reaction time corresponding to the time spent to choose each category for each trial, importance, and difficulty represents the ratings given by the participants based on their subjective assessment of the importance and difficulty of each category. We calculated the weights for each participant then took the average among the participants and tested if the weights differed from 0 to calculate the t-test for the weights.

### Factors affecting confidence during choice

We applied a linear mixed-effects regression to investigate the factors influencing confidence according to equation (4).

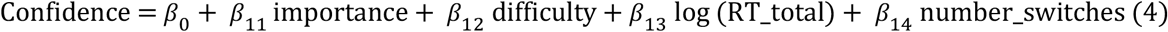

where importance is the average rating of importance in three categories based on the participants’ ratings, difficulty is the average rating of difficulty in the same three categories based on the participants’ ratings, RT_total is the reaction time of each trial, and number switches are the number of gaze switches across and within categories in each trial. We fitted the model participant by participant and then tested the betas which were significantly different from zero across all participants.

### Data availability

The data and analysis scripts have been uploaded to a GitHub repository, which can be accessed here. https://github.com/fatmaalzahraaaboalasaad/PlanningProject/tree/main

There are 45 different scenarios and stimuli used in the experiment. Some images are not licensed for sharing in the paper, but they can be provided upon request.

