## supplementary for "Prioritization of decisions by importance and difficulty in human planning"

### Supplementary materials

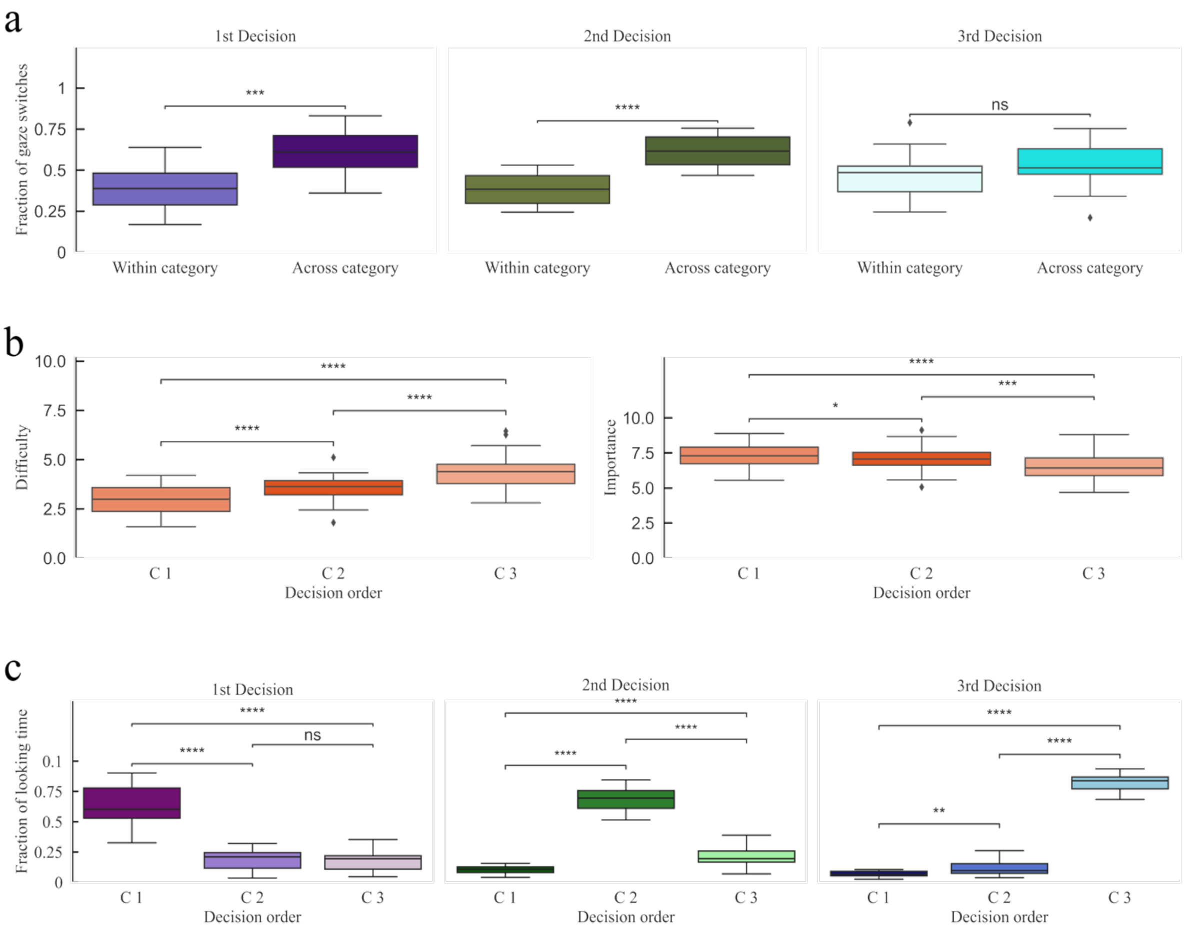


**Supplementary Figure S1**: Prioritization of decision trees and how humans build them. (**a**) Gaze switching between options. The fraction of within and across-category gaze switches are displayed and analyzed separately for 3-time windows in the decision phase indicated at the top. Boxplots correspond to the mean values obtained for all participants. Results of paired t-tests are displayed according to p-values (‘ns’; P > 0.05, ‘***’:P < 0.001,’****’: P < 0.0001**). (b)** The reported importance and difficulty ratings across categories. Categories are labeled according to the decision order of the trial. Boxplots were calculated over the means of all participants. Results of pairwise comparisons are displayed based on adjusted p-values (‘*’: P < 0.05, ‘***’: P < 0.001,’****’: P < 0.0001). (c) Fraction of looking time per category from stimulus onset until the first decision. Categories are labeled according to the decision order of the trial. Boxplots were calculated over the means of all participants. Results of pairwise comparisons are displayed based on adjusted p-values (‘ns’:P > 0.05, ‘****’: P < 0.0001).

| N_Scenario | written_scenario | Categories |
|---|---|---|
| **1.** | Plan a barbecue | **Sauce**<br>Aioli<br>Spicy sauce<br>Ketchup<br>**Side**<br>Salat<br>Bruschetta<br>Potatoes<br>**Food**<br>Sausage<br>Steak<br>Vegetable |
| **2.** | Plan how to study for an exam | **Location**<br>Library<br>Café<br>At home<br>**Method**<br>Text highlighting<br>Summary<br>Flashcards<br>**People**<br>Alone<br>Study partner<br>Study group |
| **3.** | Plan which ingredients you want to use for cooking dinner today | **Starchy food**<br>Potato<br>Pasta<br>Rice |

| | | |
|---|---|---|
| | | **Vegetables**<br>Tomato<br>Paprika<br>Broccoli<br>**Protein source**<br>Chicken<br>Tofu<br>Salmon |
| **4.** | Plan a movie night with a friend | **Genre**<br>Crime<br>Superheroes<br>Fantasy<br>**Place**<br>Car Cinema<br>Cinema<br>Home Cinema<br>**Snacks**<br>Nachos<br>Popcorn<br>Ice-cream |
| **5.** | Plan a game night at your place | **Drink**<br>Cocktail<br>Sparkling wine<br>Beer<br>**Participants**<br>3<br>4 |

| | | |
|---|---|---|
| | | 2<br>**Game**<br>PlayStation<br>Card game<br>Board Games |
| **6.** | Plan a birthday party for your younger self when you turn eight | **Theme**<br>Kings and Queens<br>Magicians<br>Astronauts<br>**Decoration**<br>Confetti<br>Balloons<br>Steamer<br>**Activity**<br>Sack race<br>Water fight<br>Ball pit |
| **7.** | Plan your wedding | **Venue**<br>Beach<br>Garden<br>Indoor<br>**Cake**<br>Fruit<br>White<br>Mini cakes<br>**Flowers** |

| | | |
|---|---|---|
| | | Peonies<br>Roses<br>Lilies |
| **8.** | Plan your Sunday breakfast | **Drink**<br>Tea<br>Coffee<br>Juice<br>**Food**<br>Pancakes<br>Porridge<br>Toast with Egg<br>**Fruit**<br>Strawberry<br>Blueberry<br>Raspberry |
| **9.** | Plan the interior of your living-room | **Paint**<br>color Green<br>Blue<br>Red<br>**Lamp**<br>Gold metal<br>Black geometric<br>Black classical<br>**Sofa**<br>Brown leather |

| | | |
|---|---|---|
| | | Green velvet<br>White leather |
| **10.** | Plan a picnic with friends | **Seating**<br>Blanket<br>Bench<br>Chair<br>**Snacks**<br>Fruit platter<br>Cheese Platter<br>Ham and melon<br>**Sport**<br>Football<br>Badminton<br>Frisbee |
| **11.** | Plan a restaurant visit with your family on the weekend | **Time**<br>12:30<br>15:30<br>20:30<br>**Cuisine**<br>Indian<br>Italian<br>Japanese<br>**Price**<br>$$$<br>$ |

| | | |
|---|---|---|
| | | $$ |
| **12.** | Plan a hike in nature on a free day | **Location**<br>Woods<br>Mountains<br>Cliffs<br>**Duration**<br>1h<br>3h<br>2h<br>**Snack**<br>Dried fruit<br>Nuts<br>Apple |
| **13.** | Plan the ingredients to mix yourself a cocktail | **Herb**<br>Mint<br>Cinnamon<br>Ginger<br>**Fruit**<br>Lemon<br>Lime<br>Orange<br>**Alcohol**<br>Rum<br>Champagne<br>Martini |

| | | |
|---|---|---|
| **14.** | Plan a romantic date this weekend | **Present**<br>Wine<br>Chocolate<br>Flowers<br>**Outfit**<br>Casual<br>Elegant<br>Sporty<br>**activity**<br>Cocking<br>Pottery<br>Gallery visits |
| **15.** | Plan your own Pizza for dinner | **Spice**<br>Garlic<br>Basil<br>Chili<br>**Sauce**<br>Tomato<br>Cheese<br>Pesto<br>**Topping**<br>Olives<br>Pepperoni<br>Ham |

| | | |
|---|---|---|
| **16.** | Plan your own Pasta for lunch | **Pasta type**<br>Spaghetti<br>Penne<br>Macaroni<br>**Sauce**<br>Cream<br>Tomato<br>Meat sauce<br>**Topping**<br>Bacon<br>Aubergine<br>Parmesan |
| **17.** | Plan a delicious cake for the birthday of a friend | **Dough**<br>Orange<br>Chocolate<br>Plain<br>**Frosting**<br>Chocolate<br>Strawberry<br>White<br>**Toppings**<br>Sprinkles<br>Almonds<br>Chocolate sprinkles |
| **18.** | Plan a day in Barcelona with a friend who is new to the city | **Building** |

| | | |
|---|---|---|
| | | Sagrada<br>Casa Batllo<br>Park Guell<br>**Shoes**<br>Flip flops<br>Sneaker<br>Sandals<br>**Food**<br>Paella<br>Patatas bravas<br>Calamari |
| **19.** | Plan your Saturday night | **Place**<br>Bar<br>Dance club<br>Fair<br>**Participants**<br>Huge group<br>Date<br>Best friend<br>**Jacket**<br>Leather<br>Jeans<br>Light fabric |
| **20.** | Plan how you want your garden to look like | **Tree**<br>Flower tree |

| | | |
|---|---|---|
| | | Fruit tree<br>Large tree<br>**Plants**<br>Vegetables<br>pink flowers<br>yellow flowers<br>**Furniture**<br>Chairs<br>Rocking chair<br>Sofa |
| **21.** | Plan a tea party with your friends | **Tea**<br>Green<br>Black tea<br>Red fruits<br>**Setting**<br>Garden<br>Teahouse Hotel<br>**Sweets**<br>Brownies<br>Cupcakes<br>Cookies |
| **22.** | Plan your sandwich for lunch | **Bread**<br>9-Grain Wheat<br>Italian<br>Flatbread |

| | | |
|---|---|---|
| | | **Vegetables**<br>Cucumbers<br>Lettuce<br>Tomatoes<br>**Cheese**<br>Brie<br>Mozzarella<br>Cheddar |
| **23.** | Plan a beach day with your friends | **Shirt**<br>Hawaii shirt<br>T-shirt blue<br>White top<br>**Activity**<br>Stand up paddle<br>Inflatable<br>Snorkel<br>**Sun protection**<br>Hat<br>Beach umbrella<br>Beach Tent |
| **24.** | Plan a city trip | **Season**<br>Summer<br>Autumn<br>Winter<br>**City** |

| | | |
|---|---|---|
| | | London<br>Paris<br>Amsterdam<br>**Luggage**<br>Backpack<br>Suitcase<br>Bag |
| **25.** | Plan your Monday morning on a workday | **Transportation**<br>Walking<br>Bus<br>Metro<br>**Coffee place**<br>To go<br>In the kitchen<br>In bed<br>**Activity**<br>Newspaper<br>Mediate<br>Work out |
| **26.** | Plan how you want to read a book in your free time | **Seating**<br>Armchair<br>Park<br>bench<br>**Café**<br>Medium Tablet |

| | | |
|---|---|---|
| | | Modern book<br>Old book<br>**Genre**<br>Adventure<br>Fantasy<br>Detective |
| **27.** | Plan how to wash one round of your laundry | **Laundry**<br>Dark<br>Light<br>Color<br>**Drying**<br>Dryer<br>Clothesline<br>Drying rack<br>**Detergent**<br>Liquid<br>Powder<br>Tab |
| **28.** | Plan what festival to visit with your friends | **Music**<br>Rock-band<br>Reggae<br>DJ<br>**Location**<br>City<br>Lake |

| | | |
|---|---|---|
| | | Desert<br>**Event**<br>Fireworks<br>Air colors<br>Laser show |
| **29.** | Plan a one-week camping trip in Spain | **Tent**<br>Big tent<br>Small tent<br>Car tent<br>**Sleeping**<br>Hammock<br>Inflatable Mattress<br>Sleep mat<br>**Food**<br>Barbecue<br>Gas stove<br>Campfire cooking |
| **30.** | Plan a day in the mountains | **Activity**<br>ski<br>Snowboard<br>Sleight<br>**Transport**<br>Chairlift<br>Gondola<br>Platter lift |

| | | |
|---|---|---|
| | | **Hill**<br>Not steep<br>Really steep<br>Medium steep |
| **31.** | Plan your sport work out for tomorrow | **Sport class**<br>Spinning<br>Yoga<br>Bouldering<br>**Pants**<br>Short<br>Jogger<br>Tights<br>**Shake**<br>Protein<br>Green<br>Carrot |
| **32.** | Plan your order at a restaurant | **Wine**<br>Red<br>White<br>Rose<br>**Meal**<br>Ratatouille<br>Chicken<br>Pasta<br>**Dessert** |

| | | |
|---|---|---|
| | | Creme brulee<br>Tiramisu<br>Strawberry cake |
| **33.** | Plan how you want to relax after a workday | **Snack**<br>Ice cream<br>Chips<br>Fruits<br>**Entertainment**<br>Amazon video<br>Netflix<br>You tube<br>**Location**<br>Balcony<br>Sofa<br>Bed |
| **34.** | Plan the items to decorate your Christmas tree | **Christmas balls**<br>Red<br>Shiny<br>Purple<br>**Candles**<br>White<br>Gold<br>Red<br>**Decoration**<br>Candy canes |

| | | |
|---|---|---|
| | | Tree hanger<br>Animal hanger |
| **35.** | Plan what car you want to buy | **Color**<br>Black<br>Yellow<br>Red<br>**Model**<br>Bulli<br>Sports car<br>Old-timer<br>**Seats**<br>Brown leather<br>Black leather<br>Light fabric |
| **36.** | Plan how you want to do your grocery shopping | **Store**<br>Organic<br>Market<br>Supermarket<br>**Food type**<br>Carbohydrates<br>Meat, fish, cheese<br>Vegetables, fruits<br>**Bag**<br>Plastic<br>Backpack |

| | | |
|---|---|---|
| | | Trolley |
| **37.** | Plan your shopping for new clothing | **Quantity**<br>1<br>Many<br>2<br>**Shop**<br>Second hand<br>Mall<br>**Shopping street**<br>Item Jumper<br>Jeans<br>Shirts |
| **38.** | Plan your cleaning routine | **Soap**<br>Lavender soap<br>Honey soap<br>Liquid soap<br>**Time**<br>Night<br>Morning<br>Evening<br>**Place**<br>Bathtub<br>Shower<br>Sink |

| | | |
|---|---|---|
| **39.** | Plan how you want to go for a run | **Music**<br>Earphones<br>Headphones<br>No music<br>**Location**<br>Beach<br>Indoor<br>Park<br>**Time**<br>15<br>30<br>45 |
| **40.** | Plan what kind of watch you want to buy | |
| **41.** | Plan to see a sports event | **Sport**<br>Basketball<br>Tennis<br>Soccer<br>**Watching place**<br>Arena seating<br>Arena standing<br>Livingroom<br>**Fan article**<br>Scarf<br>Cap<br>Shirt |

| | | |
|---|---|---|
| **42.** | Plan how you want to have dinner tonight | **Food source**<br>Cooking<br>Ordering<br>Going for dinner<br>**Company**<br>Alone<br>Two people<br>Three people<br>**Portion size**<br>Small<br>Medium<br>Large |
| **43.** | Plan how to draw a painting | |
| **44.** | Plan your dream house | **Material**<br>Brick<br>Concrete<br>Wood<br>**Property**<br>Coastline<br>City<br>Countryside<br>**Windows**<br>Window front<br>Vintage<br>Round |

| | | |
|---|---|---|
| **45.** | Plan a party at your place | **Music**<br>90s<br>2000<br>Jazz<br>**Activity**<br>Beer-pong<br>Dancing<br>Karaoke<br>Drinks<br>**Punch**<br>Beer<br>Wine |

**Supplementary Table ST1:** Planning scenarios and stimuli. Each scenario consists of three categories and each category contains three items, with the category type in bold.
